# When are inter-individual brain-behavior correlations informative?

**DOI:** 10.1101/036772

**Authors:** Maël Lebreton, Stefano Palminteri

## Abstract

Characterizing inter-individual differences induced by clinical and social factors constitutes one of the most promising applications of neuroimaging. Paving the way for such applications, neuroimaging studies often report between-group differences in “activations” or correlations between such “activations” and individual traits. Here we raise cautionary warnings about some of those inter-individual analytic strategies. These warnings become critical when measures of “activations” are unstandardized coefficients of regressions between BOLD signal and individual behavior.

First, using simple algebraic derivations, we show how inter-individual differences results can spuriously arise from neglecting the statistical relationships which link the ranges of individual BOLD activation and of recorded behavior. We also demonstrate how apparently contradictory results and interpretations may simply arise from the interaction of this scaling issue and the pre-processing of the behavioral variables. Second, using computational simulations, we illustrate how this issue percolates the booming field of model-based fMRI. Finally, we outline a set of recommendations, which might prove useful for researcher and reviewers confronted with questions involving inter-individual differences in neuroimaging.

**Author Summary:** Characterizing inter-individual differences induced by clinical and societal factors constitutes one of the most promising applications of neuroimaging. Paving the way for such applications, an increasing fraction of neuroimaging studies reports between-group differences in “activations” or correlations between “activations” and individual traits. In this manuscript, we focus on the typical analytical strategies employed in studies investigating how differences in behavior between individuals or groups of individuals are translated to differences of activations in specific brain regions. We notably question whether they are suitable to support inferences and claims about the neural underpinnings of differential cognition. Our core results demonstrate that typical inter-individual results can spuriously arise by overlooking plausible statistical relationships that link the ranges of individual BOLD activation with the ranges of produced behavior. We argue that these results challenge current classical interpretations of inter-individual results. Highlighting the methodological and theoretical gaps regarding the analysis and interpretation of inter-individual differences is fundamental to fulfilling the promises of neuroimaging.

## INTRODUCTION

“*There is very little difference between one man and another; but what little there is, is very important*”^i^. Understanding the average — *typical* — brain and how individuals differ from one another constitute the two complementary goals of cognitive neuroscience. Inter-individual differences thereby constitute a source of statistical noise when considering the typical brain, but may also represent the very object of interest (Braver et al., 2010; Gabrieli et al., 2015). Understanding how inter-individual differences in brain functions relate to inter-individual differences in behavior lies at the heart of the most promising applications of neuroimaging, such as improving neuropsychiatric diagnostics or refining individual social or cognitive traits characterization (Camerer et al., 2005; Mennes et al., 2011; Wiecki et al., 2015).

So far, one of the dominant strategies employed to investigate these research questions has leveraged task-evoked activations (see Gabrieli et al., 2015; Kanai and Rees, 2011 for alternative strategies). The intuitive analytical strategy consists of estimating individual brain activations for a given contrast in a region of interest (ROI) and testing the statistical associations between these “activations” and external heterogeneity factors. Depending on the research questions, external heterogeneity factors can be scores from clinical tests (e.g. typical symptom severity scales), psycho-socio-economic measures, i.e., “traits” (e.g. the socio-economic status), summary statistics derived from behavioral recordings, i.e., “task performance” (e.g. average accuracy, reaction times), or computational model parameters (e.g. learning rates, statistical decision thresholds) (**Figure 1.A** and **1.B**). Overall, these analyses are referred to as inter-individual brain-behavior differences (IIBBD).

**Figure 1:**
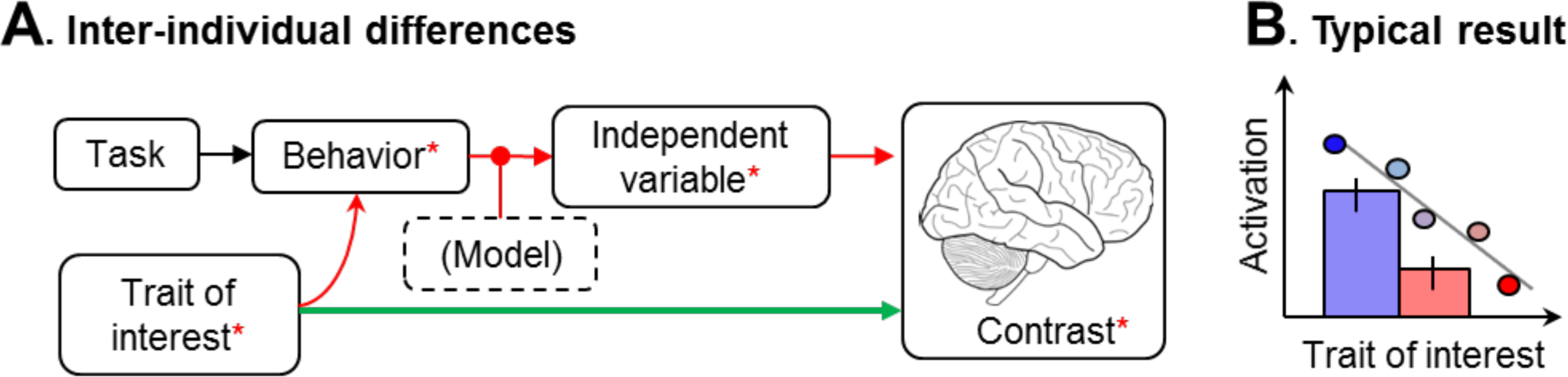
Inter-individual brain-behavior differences (IIBBD) **A**. Inter-mdividual bram-behavior differences (IIBBD) are paramount in virtually all fMRI designs, regardless ofwhether they address these chfferences explicitly or not. They are typically assessed by linking individual fMRI contrast values with a trait of mterest (green path). Problems can arise when the trait of mterest also generates chfferences in the vanance of the behavioral vanables, through complex statistical dependencies (red path). **B**. Typical results illustramg IIBBD, as they are reported in the literature, in the form of categorical contrast between group of subjects (bars±s.e.m) or group level linear correlation with a conmuous trait (dots and regression line).

In the present paper, we raise two fundamental and related questions about the standard IIBBD analytical strategy.

First, we question the *non-independence* between subject (or ffist-) ‐and group (or second-) ‐level statistics. Although significant IIBBD are often presumed statistically independent from classical average population analyses (random-effect), the validity of this assumption is unclear. This non-independence issue could notably become problematic when IIBBD analyses are used to make inferences, e.g., to select ROIs within a network of activations.

Second, we cast doubts on the standard *inteipretations* inter-individual differences. Currently, most correlations are interpreted in terms of individual neural resource mobilized to complete the task. However as noted by other authors (Poldrack, 2015; Yarkoni and Braver, 2010), these interpretations are with little prior rationale and little consistency. It is paradigmatic in the example of executive control literature, where positive correlations between prefrontal activations and performance measures are interpreted as an effective increase in cognitive control mobilization, whereas negative correlations are interpreted as an increase in neural efficiency (Poldrack, 2015; Yarkoni and Braver, 2010).

In the following paper, we explore these two issues in the case of event-related parametric fMRI designs, where individual activations are derived from individual behavior. After recalling critical differences between different metrics used as activation measures, we focus on unstandardized regression coefficients, i.e., *betas*. In the context of classical or model-based task-related fMRI, we argue that neglecting the statistical dependence of regression coefficients on how the brain signal scales with the behavior, makes IIBBD at best uninformative, and at worse misleading.

## RESULTS-PART 1. Introducing the scaling issue in inter-individual brain-behavior analyses

### fMRI analysis background

The classical fMRI analysis scheme relies on the general linear model (GLM) framework (Friston et al., 1994), and follows a *multi-level summary statistics* approach (Beckmann et al., 2003; Friston et al., 2005; Holmes and Friston, 1998; Woolrich et al., 2004; Worsley et al., 2002), to approximate mixed-effects designs (Beckmann et al., 2003; Friston et al., 2005; Mumford and Nichols, 2009). This approach is briefly summarized as follows: in a first step, the linear relation between the time series of BOLD signal and the time series of the different explanatory variables is assessed at the individual level. For each individual *k*, this entails designing a first-level GLM:

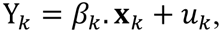

where ***Y***_*k*_ is the BOLD time-series, ***x***_*k*_ contains the explanatory variables time-series, *u*_*k*_ is a Gaussian noise, and *β*_*k*_ is the vector of the unstandardized linear coefficient of regressions to be estimated. First-level summary statistics, i.e., estimated individual betas 
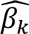
 or contrasts of betas, are then used in a population level analysis. For studies aiming at mapping a cognitive function in the typical brain, this second-level analysis is usually a random effect, i.e., a “one-sample t-test”. However, second-level analyses can also aim at investigating differences in activations between different categories subject (e.g. pathological or non-pathological sub-populations), or across a continuum of subjects (e.g. following individual “traits”). These IIBBD analyses can respectively be implemented with “two-sample t-tests” or ANOVAs for the categorical case, and with second-level “multiple regression” or “correlations” for the continuous case (**Figure.1.B.**).

### Measures of activations

In order to address issues in IIBBD analyses, we must first define what typical individual measures of “activations” are. As recalled in the previous paragraph, the most common measure is the first-level unstandardized coefficient of regression, or betas 
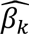
. For each independent variable **x**_*i*_ (Cohen et al., 2013):

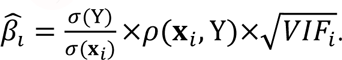

Here, *ρ*(**X**_*i*_,**Y**) is the semi-partial correlation between **x**_*i*_ and **Y**, and ***VIF***_*i*_ is the Variance Inflation Factor, which quantifies *β̂*_*L*_ over-estimation due to multi-collinearity issues (i.e. due to the correlations between **x**_*i*_ and the other independent variables **x**_*j,j*≠*i*_). Hence, a fundamental property of *β̂*_*L*_ is that their value is proportional to the linear dependency between the dependent and the independent variable *ρ*(**X**_*i*_,Y), but also to the ratio of their standard deviation 
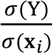
—i.e, to the scaling of those variables.

One can also compute *t̂*_*L*_, the *Student t-statistic of β̂*_*L*_, relative to the null hypothesis *β̂*_*L*_ = 0. Assuming a Gaussian noise *u* in (1), we have

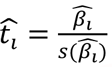

Here, s(*β̂*_*l*_) is the standard error of the estimate *β̂*_*l*_,

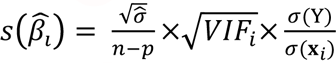

where σ^2^ quantifies the variance of the errors of the regression model, *n* is the sample size, and *p* is the number of coefficients in the model including intercept. Hence, one can show that

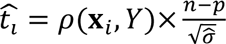

Therefore, as opposed to *β̂*_*l*_,*t̂*_*i*_ depends on the linear dependency between the dependent and the independent variable and on the overall quality of the regression model, but not on the scaling of the variables. *t̂*_*l*_ follow a Student’s t-distribution with (*n* – *p*) degrees of freedom, from which the P-value corresponding to the null hypothesis *β̂*_*l*_ = 0 can be computed.

*Z*-*values* ***Ẑ***_*l*_, which are sometimes preferred to *t*-*values* because they are independent of the number of degrees-of-freedom, are typically re-computed from these P-values.

Up to this point, one can already stress that the term “activation” for a specific contrast and a specific individual is misleading given that different measures (*β̂*,*t̂*,*Ẑ* are available, which represent different quantities and bear different interpretations. It seems critical, for future studies of IIBBD, to clarify the question they are addressing and make appropriate choices, descriptions and interpretations of their activation measure: while *t̂* or *Ẑ* values seem appropriate to investigate whether an ROI encodes a variable with a different *reliability* in different subjects, *β̂* values additionally carry information about *differences in scaling* between the variable and the brain signal in different subjects.

### Scaling laws and brain-behavior correlations

Critically, most IIBBD analyses rely on *unstandardized* first-level betas 
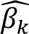
, which as recalled earlier, depend on the ratio of the standard deviation of the BOLD σ(Y_*k*_) to the standard deviation of the experimental factor σ(x_*k*_). We refer to the statistical relationships between σ(Y_*k*_) and σ(x_*k*_) as *scaling laws*. Although, as we will demonstrate in this paper, understanding how the brain signal scales with the behavior is fundamental to draw theories and perform IIBBD analyses, very little is known about how the BOLD signal range varies with different ranges of stimulations or of produced behavior. We consider two hypothetical, though neuro-biologically plausible, scaling laws between a behavioral measure and the BOLD signal: the *proportional* and the normalization hypotheses. Without loss of generality, we consider those scaling laws in an idealized situation, where the BOLD signal in a brain region ***Y***_*k*_ encodes a parametric measure of interest ***x***_*k*_ whose range σ(x_*k*_) varies between individuals *k* (see **Figure.1. A** and **B**, and **Experimental procedures** for formal definitions). Under the *proportional hypothesis*, the individual relationship between the brain signal and the produced behavior captured by the 
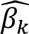
 is kept constant across individual, notwithstanding random variations: this is the implicit assumption underlying random-effect analyses in neuroimaging. As a consequence 
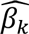
 does not correlates with σ(**x**_*k*_)(**Figure 2.A.a**). In other words, under the proportional hypothesis, *behavioral differences are reflected by underlying differences in brain activity, but not by differences in* 
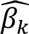
. Under the *normalization* hypothesis, the BOLD signal encodes the behavioral variable ***x***_*k*_ on an identical scale across individuals. Then, inter-individual or between-group *differences* in 
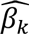
 derive from differences in behavior σ(x_*k*_) (Figure 2.B.a). Therefore, under the normalization hypothesis, *behavioral differences are not reflected by underlying differences in brain activity, but by differences* 
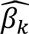
.

**Figure 2:**
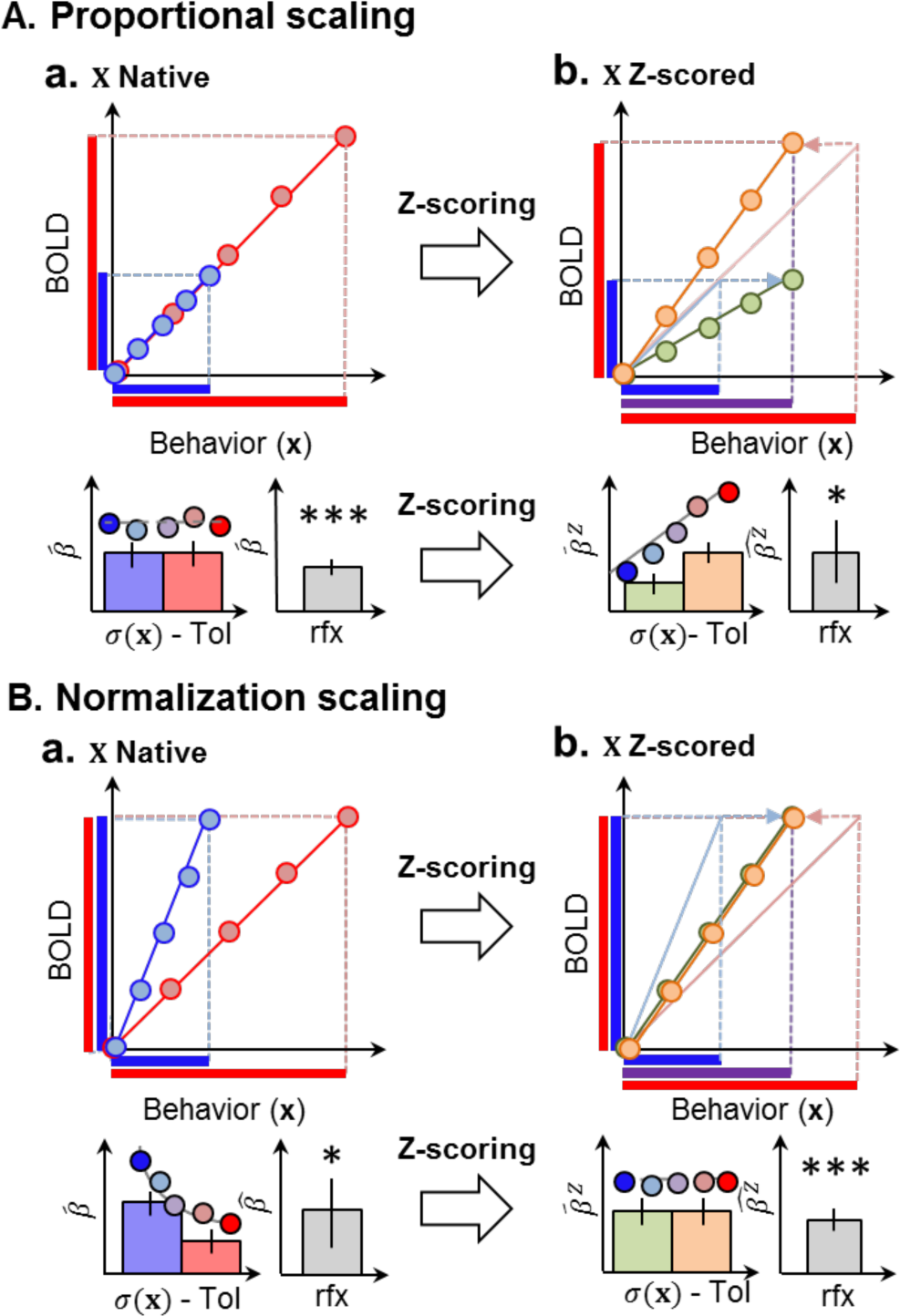
The relationship between scaling laws and second-level statistics. The panels describe the impact of mter-mchvidual differences in the standard deviahon *σ*(**x**) of the behavioral variable X, under different scaling-laws hypotheses ‐proporhonal (**A**) and normalizahon (**B**), and different preprocessmg ‐nahve (**a**) or Z-scored variable (**b**). Each sub panel contams three graphs. On the left we illustrate how **x** is related to the BOLD signal in two mdividuals wifh different Lmtial *σ*(**x**) (blue vs. red or green vs. orange). The mdividual unstandardized coefficients of regression *β̂*, corresponds the slope of the correspondmg lmes. On the upper right comer, we illustrate the stahstical relahons between mdividual bram achvahons and *σ*(**x**) (presumably lmked to the trait of mterest Tol) as a between-group analysis (histograms) or conhnuous mter-mdividual correlahon (dots). On the bottom-right comer we illustrate the consequences for the significance of second-level random-effect analysis (*: lower significance, vs. ***: higher significance). *β̂* and 
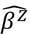
 respechvely refer to fMRI unstandardized coefficients of regressions computed with a nahve scalmg or an mdividual Z-scoring of the paramemc regressor **x**.

### The consequence of variable normalization

To circumvent the scaling issue, one might be tempted to neutralize inter-individual differences by normalizing −Z-transforming‐ the behavioral variable of interest. However, under the *proportional* hypothesis, Z-scoring induces a monotonic relationship between the “activation” 
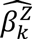
 and the standard deviation of the behavior σ(x_*k*_), which was absent before Z-scoring (**Figure 2. A.b; Experimental procedures**). Conversely, under the *normalization* hypothesis, Z-scoring cancels the monotonic relationship between 
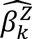
 and σ(x_*k*_), removing the correlation between the activation and the behavioral variable of interest (**Figure 2.B.b**). These derivations critically demonstrate that the interpretation of typical IIBBD results largely depends on an interaction between the pre-processing of the behavioral variable and the underlying scaling hypothesis. The interaction between the scaling hypothesis and the Z-scoring of the behavioral variable also affects the results of second-level random-effect analyses (i.e. one sample t-test on the 
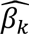
): under the proportional hypothesis, Z-scoring ***x***_*k*_ introduces a systematic variance in the 
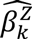
 due to the dependence of 
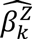
 to σ(x_*k*_), which can decrease the significance of the random-effect model (see **Figure 2.A.a** vs **2.A.b**); conversely, under the normalization hypothesis, Z-scoring ***x***_*k*_ erases the variance initially due to the difference in σ(x_*k*_), therefore increasing the significance of random-effect model (see **Figure 2.B.a** vs **2.B.b**). Thus, systematic investigation of brain-behavior scaling laws may provide precious information about how to best pre-process variables of interest to perform IIBBD analyses.

### Ignoring scaling laws constitutes a challenge for interpreting differences in activation

These derivations carry an important message: in the context of a parametric design, the interpretations of inter-individual differences in 
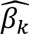
 should be made with caution. For instance, let's assume that the activity in a region of interest (e.g. the hippocampus) ***Y***_*k*_ is causally responsible for a behavioral measure ***x***_*k*_ (e.g. levels of memory strength), and that a heterogeneity factor (e.g. score on a clinical test for Alzheimer's disease) causes changes in *Y*_*k*_, inducing proportional changes in ***x***_*k*_. In the proportional context, comparisons between 
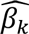
 in the ROI might be inconclusive, potentially misleading to false-negative conclusions about the role of the ROI in inter-individual differences (**Figure 3.A**).

A second implication from our derivations is that any heterogeneity factor which modulates-or correlates with-individual behavior σ(x_*k*_) is spuriously associated with individual activations measured by 
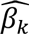
. Critically, the directionality of this statistical association is somewhat unpredictable, given that it depends on interactions between the ‐typically unknown-underlying scaling laws and the processing of the behavioral variable.

A third implication is that comparing parametric 
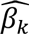
 in the presence of behavioral differences is not trivial. This not only covers neuroimaging experimental designs where researchers are investigating inter-individual differences, but also extends to within-subject between-sessions designs. Critical cases arise when behavior and BOLD activities are recorded in the same individuals but in different sessions, where σ(x_*k*_) is susceptible to be modulated. In that case, statistical tests based on the contrast of 
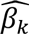
 between sessions are vulnerable to scaling issues: the interpretation of the result depends on the underlying scaling laws and the processing of the behavioral variable (e.g. between-sessions Z-Scoring). This cautionary message applies, for example, to typical experimental designs investigating the effects of a pharmacological manipulation, a stimulation protocol (e.g. trans-cranial magnetic stimulation), or a contextual effect on a behavior-related activation.

Finally, the critical implication of this demonstration is that IIBBD analyses might be not be as independent from the average population analyses as initially thought. In other words, in a ROI significantly encoding a variable at the population level, individual 
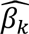
 are *de facto* linked to σ(x_*k*_), in a way that depends on the underlying scaling law. This is a fundamental issue when such analyzes are used for inferences, e.g., when used as a selection criteria to highlight the role of a specific ROI in a larger network of activation.

**Figure 3:**
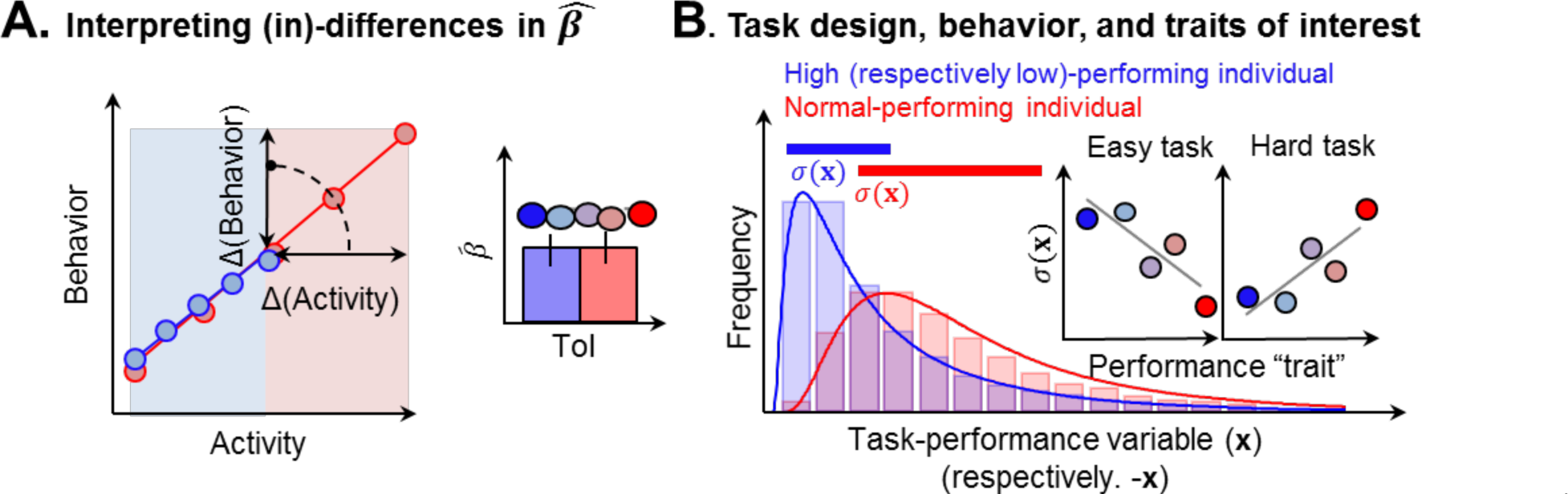
Interpreting differences in unstandardized □. **A.** Consider a bram region, which causally and proporhonally causes a behavior (i.e. the more achvahon, the higher the behavioral variable, withm and across subjects). In case a trait of mterest (e.g. pathology) directly impacts the range of achvahon of this region (e.g. due to degeneration), this cannot be assessed/detected by chfferences in *β̂* (right, mset). Msunderstandmg the significahon of *β̂* and ignormg scalmg laws can lead to erroneous negahve conclusions. B. Lmkmg task design, behavior, and traits of mterest. The red/blue histograms depict the chsmbution of a behavioral variable (reaction time, decision value, confidence) in two individual with variable performance. If the task is easy (respectively difficult), the high (respectively low)-performmg mdividual can exhibit a ceilmg (respectively floor) effect on performance. This creates statistical dependencies between the performance trait performance) and the standard deviation (σ(**x**)) of the behavioral task-performance variable (see graphical insets). Given the dependencies between fMRI*β̂* and *σ*(**x**), this can lead to opposite correlations between “activations” ‐as measured by *β*‐, and “individual performances”, proxied by *σ*(**x**).

### The importance of task design in brain-behavior correlations

Finally, the importance of task design should not be overlooked in the interpretation of IIBBD. Consider a typical fMRI study aiming at linking differences in an individual “trait” (e.g. IQ) to neural activations, which are derived from a trial-by-trial measure of a task-related performance variable ***x***_*k*_ (e.g. confidence ratings). When the task is easy, high-performing (i.e. high-IQ) subjects exhibit ceiling performance (only high confidence ratings). Accordingly, the trait of interest (IQ) will be associated to a smaller standard deviation of the task-related variable, σ(x_*k*_). This creates a negative correlation between the trait of interest (IQ) and σ(x_*k*_). Symmetrically, when the task is difficult, low-performing (i.e. low-IQ) subjects will exhibit flooring performance (only low confidence). Accordingly, “poor performance” (and therefore low-IQ) will be associated to a smaller standard deviation of the task-related variable σ(x_*k*_). This creates a positive correlation between the trait of interest (IQ) and σ(x_*k*_). Given the dependencies between brain “activations” 
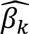
 and σ(x_*k*_), these two cases will lead to opposite inter-individual correlational results between confidence activation and IQ as a function of the overall task difficulty (**Figure 3.B**). This example clearly illustrates the potential of the scaling framework to explain inconsistent results in the cognitive neuroscience literature of IIBBD, where both positive and negative correlations between activation and performance are reported in the same brain area for a given task.

## RESULTS-PART 2-Scaling-laws and model-based fMRI

In this second section, we will investigate the consequences of the scaling issue for model-based fMRI. Model-based fMRI typically uses as dependent variables ***x***_*k*_ *latent variables*. Generally, these *latent variables* are inferred from observable behavior as follows: a computational model is fitted in order to obtain the free-parameters' values that maximize the likelihood of observing the data given the model. These free parameters are the used to calculate a trial-by-trial estimate of the *latent variables* of interest ***x***_*k*_ (O’Doherty et al., 2007). Importantly, the free-parameters can be either considered as fixed (FFX) ‐i.e. shared across individuals ‐or random-effects (RFX) ‐i.e. each subject’s parameters are drawn from a common population distribution (Daw, 2011). In the next paragraph, we discuss how choices in the parametrization of computational models can first affect σ(x_*k*_), and then percolate potential IIBBD analyses.

### Computational models and latent variable distributions

The first point of this section is that model parameters inevitably impact the distribution of the latent variable, including its standard deviation σ(x_*k*_). Consider the example of using an expected-utility model in decision-making under risk: for choice situations involving simple prospects combining potential gain(s) *g* with a probability *p* of winning, expected utility theory stipulates that agents choose the option which maximize the expected utility:

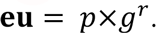

In this model *r* is the utility curvature free-parameter and captures the individual attitude toward risk (see e.g. (Bernoulli, 1954)). In order to investigate the link between the model parameter *r*, and the standard deviation of the latent variable σ(**eu**), we simulated a task where subjects are confronted with several options ‐i.e. combinations of *g* and *p*-, and computed **eu** for different and plausible values of *r*. These simulation unambiguously show that the model free-parameter *r* monotonically determines σ(**eu**), and that even moderate modulations of r (within the reasonable and plausible range [.751.25]) can cause up to a fourfold increase in σ(**eu**) (**Figure 4.A**).

**Figure 4:**
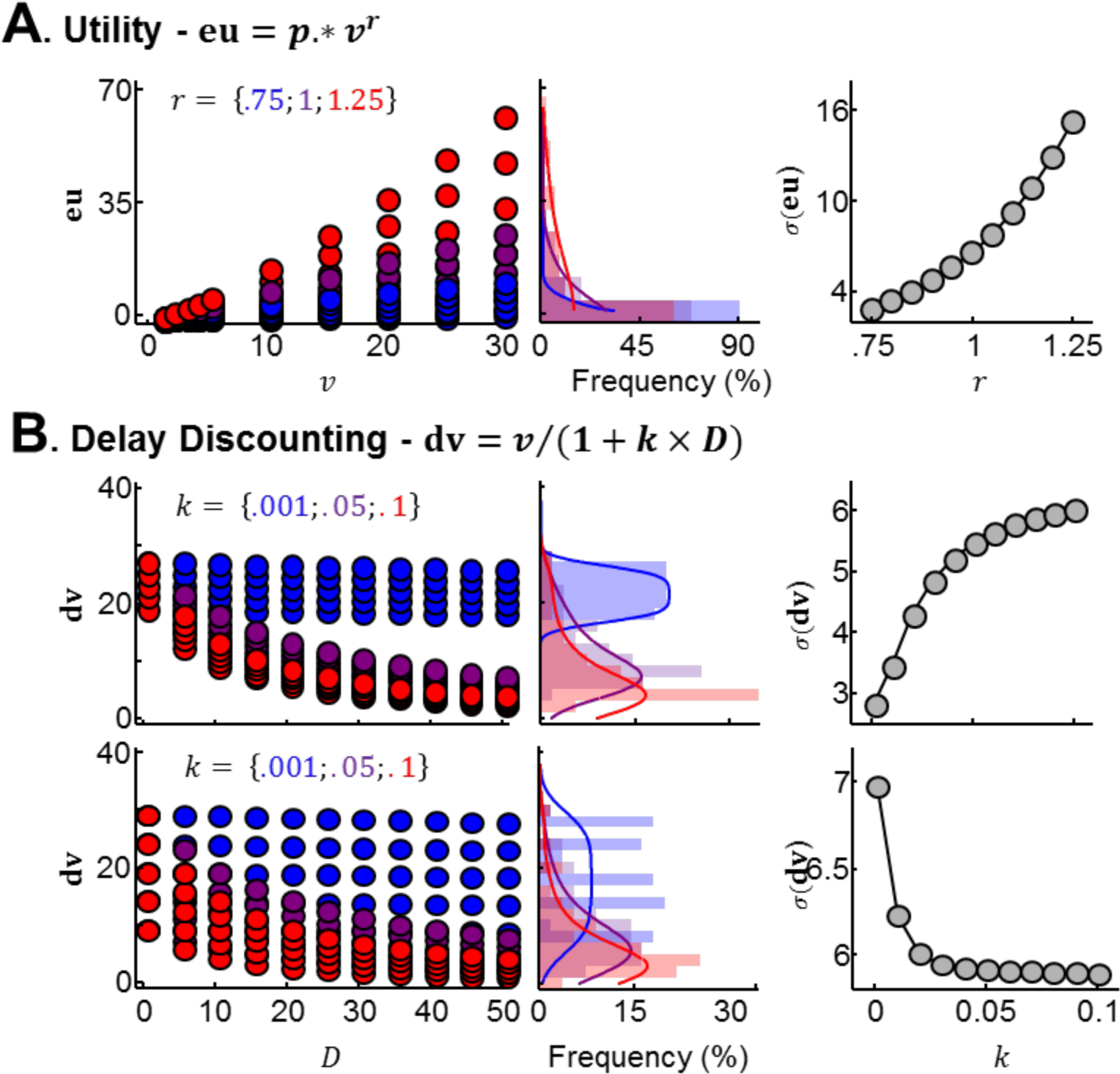
Model-based fMRI and the impact of model-parameters on inter-individual differences. **A.** Linking individual model-parameters and σ(**v**). We computed the expected utility **eu** = *p* × *g*^*r*^ of different prospects mixmg gams *g* and probabilities *p* - *g* ∈ {1:510:5:30} €, *p*   {10:20:90}%–, for different values of the uhhty curvature parameter *r*. Left: Expected uhlity of all task-smnuh for three value of *r* (blue: *r* = .75, purple: *r* = 1.0 and red: *r* = 1.25)., Middle: correspondmg histogram and scaled density funchons of expected-utility. Right: estmiated σ(**eu**) as a funchon of *r*. B. Lmhng mdividual model-parameters, task design and σ(**v**). We computed the discounted value 
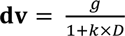
 of different prospects mixmg gams *g* and delays *D*, for 2 task designs (i.e. 2 sets of prospects). Top (task design 1): *g* ∈ {20:2:28} €,*D* ∈ {0:5:50} Bottom (task design 2): *g* ∈ {10:5:30} €,*D* ∈ {0:5:50}. Left: Discounted value of all task-shmuli for three values of the discount parameter *k* (blue: *k*= 001, purple: *k* = .04 and red: *k* = .1). Middle: correspondmg histogram and scaled density funchons of discounted values. Right: esmnated σ(**dv**) as a function of *k*.

Recall, from part 1, that “expected utility activations” ‐i.e. 
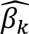
-are impacted by σ(**eu**). Hence, any inter-individual brain-behavior correlation between such activations and *r* might be spuriously driven by this statistical dependence. This raises caution and doubts toward statements such as, “*expected utility encoding in ROI X was modulated by risk-aversion*”, and this cautionary warning applies more generally to most inter-individual analyses linking model parameters and activations derived from latent variables. We advise authors to investigate how statistical relationships between model parameters and σ(x_*k*_) can affect IIBBD analyses.

### Task designs and computational models

In many cases however, the link between the model parameters and the standard deviation of the latent variable σ(x_*k*_) largely depends on the task design and the stimuli space. To illustrate this second point, we take the example of the hyperbolic delay-discounting model. This model states that in choice situations involving prospects combining future gain(s) *g* deferred by delays *D*, decision-makers are choosing the option with the highest discounted value:

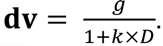

In this model *k* is the discount factor: a free-parameter that captures individual intolerance to delay (see e.g. (Ainslie and Haslam, 1992)).

In order to investigate the impact of the task design on the link between model parameters and σ(x_*k*_), we simulated two tasks where subjects are confronted with sets of options combing *g* and *D*. The only difference between the two tasks lies in the set of options, i.e., of *g* and *D*. For these two tasks, we computed the discounted value of all options for plausible values of *k*. The results of this simulation demonstrate that, depending of the option set (e.g. the task design), the model free-parameter *k* and σ(**dv**) can be either positively or negatively correlated (**Figure.4.B**). Therefore, authors should not assume or take for granted a specific directionality in the correlations between model parameters and the standard deviation of the latent variable σ(x_*k*_). Rather, we recommend to appropriately retest those correlations for every variation of an experimental setting design.

### From latent variables to choice functions

Critically, the associations between the model parameters and the standard deviation of the latent variable σ(x_*k*_) are not limited to parameters involved in the computation of the latent variable ***x***_*k*_. In the case of value-based decision-making, this means that the standard deviation of the expected value may not only be linked to parameters of the value function, but may be linked to parameters controlling the decision policy. To illustrate this point, we ran a last set of simulations, involving a simple reinforcement-learning situation, where decision-makers learn, by trial and error, to select the option associated to the higher reward rate. The Q-learning model proposes that agents learn the value of stimuli (Q-values) via a trial-by-trial iterative process: the value of the chosen option is updated at each trial with a prediction-error, which is the difference between the predicted outcome and the actual outcome *R*:

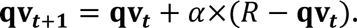

Value updating is weighted by a parameter, the learning rate *α*, which quantifies how much individuals “learn” from their errors. The decision mle between two options A and B is often implemented as a soft-max (e.g. logistic) function:

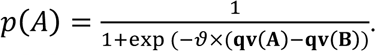

Where ϑ is a free-parameter called the temperature, and indexes the trade-off between exploration and exploitation. We first simulate learning sequences with different values of *α* and *R*, and show that these two parameters are strongly associated with σ(**qv**) (**Figure 5.A**). However, *R* is rarely set as a free-parameter, and is usually set to the outcome value, despite potential individual differences in the sensitivity to the outcome magnitude (but see (Palminteri et al., 2012) for an exception). We therefore simulated choices occurring from learning sequences with a fixed ϑ but different values of *α* and *R*, and estimated a classical (therefore “incorrect”) model, where *R* was fixed to the outcome value while *α* and ϑ were set as free-parameters. In that case, we show that both *α* and ϑ are strongly associated with σ(**qv**), despite the fact that ϑ is not a parameter governing the computation of **qv**, but rather governs the choice process (**Figure 5.B**). Consequently, while checking statistical dependencies between model parameters and the standard deviation of the latent variable σ(x_*k*_), authors should keep in mind that parameters which seem distal to the latent variable might still largely contribute to shaping its distribution.

**Figure 5:**
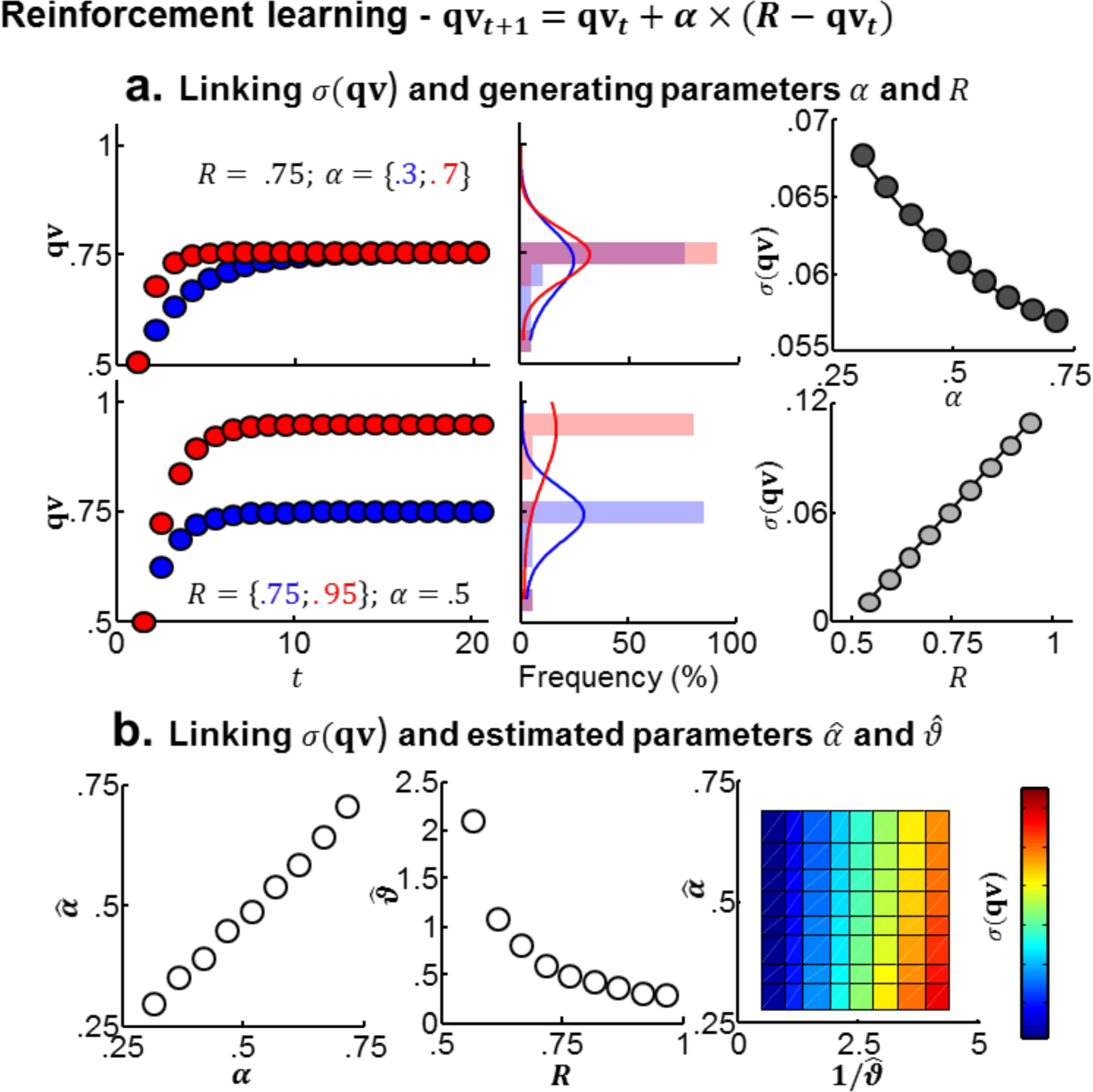
Model-based fMRI and the impact of model-parameters on inter-individual differences. **A.** Illustramg the lmk between value related model-parameters and σ(**qv**). We computed the Q-value **qv**_*t*_ + 1 = **qv**_*t*_ + *α*×(*R* − **qv**_*t*_) of a 20-trials learning sequence, for different learning rates *α* (Top, *R* = .75, and blue: *α* = .3 or red: *α* = .7) and different outcome magnitude R (Bottom; *α* = .5, and blue: *R* = .75, and red: *R* = 1) Left: Q-values as a function of the trial number. Mddle: correspondmg histogram and scaled density functions of Q-values. hght: estmiated σ(**qv**) as a function of *α* (top) or R (bottom). B. IUustrahng the lmk between non-value (i.e. decision) related model-parameters and σ(**qv**). We simulated a task, implemenmg bmary choices between a fixed-option of known value (0.5), and an option whose value had to be learned through 20 trial-and error. We generated sequences of Qv of the unknown option, with different values of *α* and R, and correspondmg stochastic choices. We then estimated the parameter of the model, but with a fixed R (=1) and a softmax (logistic) choice function, setting □ and ϑ as the model free parameters. Left: estimated *α̂* as a function of the true *α*. Mddle: estimated ϑ̂ as a function of the true *R*. hght: estimated σ(**qv**) as a function of *α̂* and 1/ϑ̂.

### The choice of the free-parameters for model-based fMRI

Although treating model free-parameters as random-effects often provide the best account of individuals' behavior as assessed by rigorous model-comparisons, a common practice is to treat them as a fixed-effect‐ i.e. using the population-level parameters-to generate the latent variables to be fed into the fMRI analysis (Daw et al., 2006; Gershman et al., 2009; Gläscher et al., 2009, 2010; O’Doherty et al., 2004; Palminteri et al., 2009, 2015; Pessiglione et al., 2008). This is often justified arguing that parameter estimates at the individual level are “noisy” and estimating them from collapsing all subjects is an efficient way to regularize them. However, if individual parameters still provide a better account of the population behavioral data according to rigorous, complexity penalizing, model-comparison procedures, then the variance modeled in the individual parameters actually captures a tme inter-individual variability. Therefore, using population-level parameters does not seem justified. As a matter of fact, the advantage of using the population-level parameters could be explained in the light of the scaling issue described above. Actually, it is worth noticing that the utilization of population-level parameters constrains σ(x_*k*_) to a unique population value, provided that individuals are given a similar input. Under the *normalization* scaling hypothesis, this can substantially increase the statistical power of subsequent second-level random effects analyses. However, under the *normalization* scaling, a more appropriate way to model brain activation would involve using individual model parameters and Z-scoring latent variables. This point further underlies the importance of better documenting scaling-laws in fMRI, so as to provide an *a priori* principled rational to preprocess independent variables of interest, in order to increase the sensitivity and replicability of model-based fMRI.

### Inter-individual correlations with model free-parameters

A consequence of the relation between the model parameters and σ(x_*k*_) is that inter-individual correlations between model free-parameters and activations 
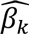
 in a ROI encoding ***x***_*k*_ may simply rely on the mathematical dependencies between the free parameters, σ(x_*k*_), and 
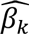
. Critically, these dependencies largely depend on the underlying scaling hypothesis. In other words, observing a significant correlation between individual learning rates and individual Q-values-related *activations* in a given area may simply reflect the fact that differences in learning rate entail differences in the Q-value standard deviation. Note that this cautionary warning can also hold when use population-level parameters for the neuroimaging analysis: using population-level parameters *de facto* underestimates the variability of the latent variable in some individuals (those whose individual-parameters would have generated a bigger σ(x_*k*_)). In those individuals and under the *normalization* hypothesis, using the population latent variable to predict brain activity leads to inflating 
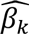
, because the (population) latent variable has to be magnified in order to match the individual’s true latent variable. The same reasoning can show that using population-level parameters deflates 
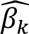
 in other individuals (those whose individual-parameters would have generated a smaller σ(**x**_*k*_)). In the end, these statistical associations provide a basis for monotonical dependencies between individual parameters values and 
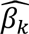
, creating spurious inter-individual brain-behavior correlation. Finally, given that the link between model-parameters and σ(x_*k*_) can reverse depending on the task design (see “**Task designs and computational models**” and Figure.4.B), these statistical dependencies can create spurious inter-individual correlations of opposite directionalities (positive or negative) between identical model-parameters and activations 
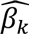
.

## DISCUSSION

### Challenges in inter-individual brain-behavior analysis

While the ability to assess and predict individual differences from neural measures-neuromarkers-has emerged as one the most promising application of fMRI in society, it is not devoid of challenges (Gabrieli et al., 2015; Wang and Krystal, 2014). In this manuscript, we explored a specific type of neuromarker: task-dependent fMRI “activations”. These are currently typically used in inter-individual brain-behavior differences (IIBBD) studies, where activations from a specific contrast in a specific ROI are correlated with a heterogeneity factor, or compared between conditions or groups of subjects. Significant differences are typically interpreted as neural underpinning of differential cognition, sometimes with opposite interpretations (efficiency vs resource mobilization) (Poldrack, 2015; Yarkoni and Braver, 2010). Critically, activations are often measured by unstandardized regression coefficients 
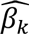
 between individual behavioral variables (**x**_*k*_) and BOLD signal (**Y**_*k*_). We claim that this poses two challenges to current IIBBD analyses. First, we question the presumed statistical independence between typical population analyses and IIBBD analyses. Second, we question of the interpretation of such analyses. These challenges rest on the simple fact that 
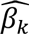
 depend on the ratio of σ(x_*k*_) to σ(Y_*k*_), making task-dependent fMRI neuromarkers dependent on the underlying *scaling* between the BOLD signal and the behavioral variable of interest. By defining two neurobiological plausible scaling laws, we demonstrated how IIBBD results can spuriously arise, and how their interpretation should be made with caution. Although we acknowledge that the present paper does not cover the full range of potential *scaling* relations between brain activation and behavior, we think that this new perspective might contribute to foster a fmitful discussion on the way to interpret and assess IIBBD in neuroimaging.

### What are scaling laws?

Scaling laws refers to the statistical relationship linking inter-individual changes in BOLD signal ***Y***_*k*_ to the inter-individual change of the related behavioral measure ***x***_*k*_. As such, scaling laws might depend of the considered behavioral variable, the experimental design and the brain region of interest.

*Stricto sensu*, scaling laws not only encompass neurobiological constraints and encoding properties of brain-behavior relationships, but are also constrained by the properties of the measures used to track brain activity, i.e., the BOLD signal. Notably, the level of interpretation of the scaling relationships may depend on the extent to which state-of-the art fMRI technics can capture inter-individual variations in the range of BOLD activations σ(Y_*k*_). Indeed, regardless of the tme underlying neurobiological coding of inter-individual behavioral differences, the *proportional* scaling hypothesis is only valid when it is possible to link inter-individual variations in the extent of the explanatory variable σ(x_*k*_) to inter-individual variations in the range of BOLD activations σ(Y_*k*_); a lack of sensitivity to relevant variations in this range of BOLD activations σ(Y_*k*_) anchors *de facto* brain-behavior analyses under the *normalization* hypothesis. Although fMRI reliability has been the focus of extensive research, this specific question has received little attention so far (Bennett and Miller, 2010). Overall, numerous factors are suspected to play a role in our ability to correctly estimate inter-individual differences in the range of BOLD activations σ(Y_*k*_), such as e.g. individual differences in vascularization or preprocessing and analytic strategies (Logothetis, 2008).

### Why does this matter?

We particularly stressed the fact that, in event-related parametric designs, differences in 
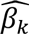
 which are used to quantify brain “activations”‐ can spuriously derive from differences in the range of the behavioral measure σ(x_*k*_), in a way that depends on the underlying scaling law. This is the case under the normalization hypothesis when neuroimaging data are analyzed with native behavioral variables **x**_*i*_, and under the proportional hypothesis, when neuroimaging data are analyzed with Z-scored behavioral variables. This adds to the fact IIBBD results might depend on the task design, and on the sensitivity of fMRI technics in capmring inter-individual variations in the range of BOLD activations σ(Y_*k*_). We therefore raise a cautionary warning about inferences derived from such analyses.

This warning is not limited to IIBBD claims, but generalizes to within-subject between-session claims: different scaling laws may have important consequences for results concerning between-session manipulations ‐such as brain stimulation or pharmacological modulations-which often also impact the behavior σ(x_*k*_), hence might spuriously impact activations 
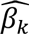
.

Our point also brings to the matter of the interpretation of IIBBD results. The proportional versus normalization scaling taxonomy is based on a formal description of the statistical dependency between the BOLD signal and the behavioral variable, rather than on a functional (over-)interpretation of such statistical quantities, like in the efficiency vs. mobilization taxonomy (Poldrack, 2015). Accurately posing and testing such hypotheses in IIBBD studies might help reconcile previous contradictory findings, and increase robusmess and reproducibility in the field.

### When does this matter?

We introduced the scaling issue in the context of event-related parametric designs, where a change of behavior σ(x_*k*_) inmitively translate in a change in activation 
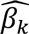
. We claim that spurious IIBBD can easily arise when traits of interests are linked to σ(x_*k*_). We also show that traits of interests impacting behavior are often linked to σ(x_*k*_) in strength and direction which non-trivially depends on task designs, making the issue of spurious correlations rather general. Critically, we also show that normalizing (Z-scoring) the behavior might not solve the problem, depending on the underlying scaling law. This issue appears to largely percolate model-based fMRI, where models parameters are used to generate latent variables and thereafter correlated with latent variable activations. In this report, we exemplified this point using models from the field of value-based decision-making, but it generalizes to different models (e.g. perceptual decision-making with drift-diffusion models).

Importantly, some critical cautionary warnings raised in this paper may be extended to fMRI categorical designs, notably when categorical events are constmcted from individual reaction times. Indeed, modelling categorical events with individual time-varying boxcars introduces an inter-individual difference in the modelling of BOLD-signal amplitude, inevitably affected by scaling issue (Grinband et al., 2008; Poldrack, 2015).

### Recommendations

We propose that a good practice before engaging in the study of fMRI IIBBD is to start documenting the statistical relationship between traits of interest (e.g. symptom severity scales) and the standard deviation of behavioral variables σ(x_*k*_) used to derive fMRI regressors 
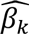
. This recommendation particularly applies to model-based fMRI, where the link between model parameters and the standard deviation of latent variables σ(x_*k*_) is sometimes not trivial. Researchers should keep in mind that even slight changes in the task design could have dramatic consequences on the strength and direction of those statistical associations. Although these recommendations are framed for IIBBD analyses, the same apply for between-session manipulations (e.g. pharmacological and other forms of brain stimulation).

Given that they can greatly influence IIBBD analysis, it is also paramount to provide detailed account of the processing of behavioral variables ***x***_*k*_ (Z-scoring or not) used for fMRI analyses.

In order to facilitate the evaluation and interpretation of IIBBD results, the term “activation” should be avoided. Multiple measures are available (*β̂*, *t̂*, *Ẑ*), which critically differs in the statistical dependency they share with dependent and independent variable. Those different measures should therefore be interpreted accurately and used to address different questions: while *t̂* or *Ẑ* values seem appropriate to investigate whether an ROI encodes a variable with a different reliability in different subjects, *β̂* values additionally carry information about differences in scaling between the variable and the brain signal in different subjects.

Ideally, the community should start explicitly investigating brain-behavior scaling laws, to avoid spurious correlations and wrong inferences in IIBBD analyses. This will entail not only deciphering the neuro-biological link between inter-individual differences in BOLD activation and behavior, but also investigating the ability of fMRI to reliably capture inter-individual variations in the range of BOLD activations. Yet, we can speculate that scaling laws differ for different cognitive functions, different brain areas, and different tasks.

Overall, in order to improve the reliability and reproducibility of IIBBD results in fMRI, rather than inmitively correlating traits and activations of interest, it is paramount to formulate clear *a priori* hypothesis about the underlying brain-behavior scaling law, to use an appropriate operationalization, and to draw accurate and cautious interpretations.

## EXPERIMENTAL PROCEDURES

### Deriving relationships between 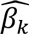 and σ(x_*k*_) under two scaling laws

We consider an idealized situation, where, in each individual *k* the BOLD signal in a brain region/voxel(Y) encodes one behavioral parametric measure of interest (**x**_*k*_) whose distribution-hence standard deviation σ(x_*k*_)-varies between individuals due to a heterogeneity source. Individual activations are capture by unstandardized regression coefficients 
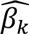
, with

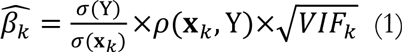

We assume that *ρ*(**X**_*k*_,Y) and 
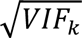
 are independent of σ(x_*k*_) and σ(Y_*k*_), i.e. that the quality of this encoding is similar across subjects and does not depend on the individual brain activation or behavioral variable or range. In other words, we set

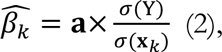

where **a** is a constant.

We can now formalize the *proportional* hypothesis by setting

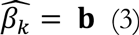

By neglecting inter-individual differences in *ρ*(**x**_*k*_,**Y**) and *VIF*_*k*_ (2) and (3) imply,

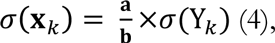

i.e. the BOLD activation scale proportionally with the behavior. Likewise, we can formalize the *normalization* hypothesis by setting

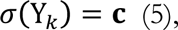

Where **c** is a constant.

In this case, (2) and (5) imply

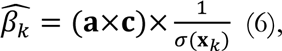

i.e., the activation summary statistics is inversely correlated with the standard deviation of the behavior.

### Scaling laws and Z-scoring

Z-transforming individually a measure of interest entails subtracting its original mean *μ*_*X,k*_ and dividing the resulting centered variable by its original standard deviation σ(x_*k*_):

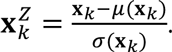

Let us assume that the two scaling hypothesis are still related to the original variables, i.e. (3) and (5) still hold, but that first level analysis are conducted with normalized variables, i.e. 
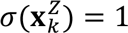
, hence

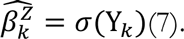

Hence, under the *proportional* hypothesis, (4) and (7) imply

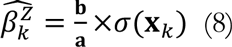

while under the *normalization* hypothesis, (5) and (7) imply

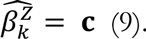

### Computational simulations

All simulations were performed in Matlab^®^

#### Linking utility parameter and **σ(eu)**

We computed the expected utility **eu** = *p* × *g*^*r*^ of different prospects mixing gains *g* and probabilities *p* − *g* ∈ {1:510:5:30} €,*p* ∈ {10:20:90}%. The standard deviation of the resulting expected utility distribution was then calculated, for different values of the utility curvature parameter *r*− *r* ∈ {.75:.05:.95}.

#### Linking discount factors, task design and **σ(dv)**

We computed the discounted value 
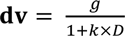
 of different prospects mixing gains *g* and delays for 2 task designs (i.e. 2 sets of prospects). Prospects from the first task design combined gains and delays from the following distributions: *g* ∈ {20:2:28} €,*D* ∈ {0:5:50}, while prospects form the second task design combined gains and delays from the following distributions: *g* ∈ {10:5:30} €,*D* ∈ {0:5:50}.

The standard deviation of the resulting discounted value distribution was then calculated, for different values of the discount parameter *k* − *k* ∈ {.001.01:.01:.10}.

#### Linking learning rates and subjective outcome to **σ**(**qv**)

We computed the Q-value **qv**_*t* + 1_ = **qv**_*t*_ + *α*×(*R* − **qv**_*t*_) of a 20-trials learning sequence, for different learning rates *α* − *α* ∈ {.5:.05:.7} and different outcome magnitude R − *R* ∈ {.55:.05:.95}.

#### Linking parameters of the decision-function to **σ**(**qv**)

We simulated a task, implementing binary choices between a fixed-option of known value (0.5), and an option whose value had to be learned through 20 trials and errors. We generated sequences of Qv of the unknown option, with different values of *α* − *α* ∈ {0.5:.05:.7}‐ and R − *R* ∈ {.55:.05:.95}.‐, and corresponding stochastic choices. For each set of parameters, we generated 50 learning-sequences. We then estimated the parameter of the model, by maximizing the likelihood of observed choices under a softmax (logistic) decision mle 
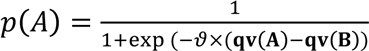
. Critically, in the estimation phase, we set a fixed R(=1), while letting *α* and ϑ vary, as the model free parameters.

## Acknowledgments

During the preparation of this work, ML was supported by an EU Marie Sklodowska-Curie Individual Fellowship (IF-2015 Grant 657904), a NWO Veni (Grant 451-15-015), a Universiteit van Amsterdam-Amsterdam Brain and Cognition Talent Grant, and acknowledge the support of the Bettencourt-Schueller Foundation. SP was supported by an EU Marie Sklodowska-Curie Individual Fellowship (PIEF-GA-2012 Grant 328822) and is currently supported by an ATIP-Avenir grant.

## Footnotes

i William James (1897). The Importance of Individuals, In *The Will Believe and Other Essays in Popular Philosophy*

